# The Implicit Assumptions of Classic Functional Responses and Their Multi-Species Extensions

**DOI:** 10.1101/2022.07.18.500336

**Authors:** Gian Marco Palamara, José A. Capitán, David Alonso

## Abstract

Functional responses are central to describe consumer-resource interactions. Defined as the per capita average feeding rate of consumers, since Holling’s seminal papers, they have been widely used in ecology. Holling’s central observation was that they often saturate as resource density increases. If the interference between consumers is strong, they also decrease with consumer density. Here we emphasize the stochastic nature of the individual feeding processes and the associated probability distributions of the different behavioral types and show how they control population-averaged feeding rates. We do so by revisiting a classic approach based on the formulation of feeding interactions in terms of individual-based reaction schemes. We highlight the common assumptions underlying the different functional forms and discover a new predator-dependent functional response that should be considered the natural extension of the Holling type II functional response when consumers interference is considered. Our work has clear implications, on the one hand, for both model selection and parameter inference from feeding experiments, and, on the other, for the use of multi-species extensions of these functional responses in population-level food-web dynamic models.

## Introduction

Functional responses measure how per capita average feeding rates of consumers respond to density of both resources and consumers. From an empirical perspective, feeding rate models are used to explain feeding experiments [1, 2, 3, 4, 5]. From a theoretical perspective, functional responses are also introduced in food-web models to explore central questions in ecology that range from ecosystem stability [6, 7] to the role of biodiversity in maintaining ecosystem functions in response to global change [8, 9]. Both applications would benefit a great deal from a careful examination of the underlying assumptions involved in the derivation of these functional forms.

Consumers undergo different processes while feeding on resources, which can be represented by chemical-like reaction schemes. To our knowledge, this approach was first used by Real [10], who re-examined the derivation of the Holling type II functional response, and derived, for the first time, the Holling type III functional response in a clear analogy to allosteric enzymatic reactions. This functional response would arise as a consequence of an increasing attacking efficiency of predators as they become exposed to higher and higher resource densities. The predator-dependent Beddington-DeAngelis [11, 12] functional response has been also framed in terms of chemical-like reaction schemes [13], although we suggest here a more consistent derivation based on a novel reaction scheme that includes both direct and resource mediated interference.

By revisiting then the derivation of these two classic functional responses (Holling type II [14, 15] and Beddington-DeAngelis [11, 12]), surprisingly, we discover that the classic form for the latter does not result from the underlying individual feeding processes in the same exact and consistent way as the former does, and, as a consequence, it should be interpreted rather phenomenologically, or, at most, as an approximation. Here, our main goal is to emphasize, on the one hand, the sometimes overlooked common assumptions underlying all these derivations, and, on the other, the inherent stochastic nature of the feeding process [16]. Our approach can be extended to more general functional responses, as long as the hypotheses underlying the feeding process can be represented by discrete events described by individual reactions, and, more generally, formalized in terms of one-step Markov processes in continuous time [17]. As an example, based on a particular hypothesis about individual processes of a consumer species feeding on multiple resources, we also derive several multi-species extensions of the Holling Type II functional response. Interestingly, we show that different hypotheses lead to different alternative extensions of the Holling type II functional response. All these multi-species extensions can be used to explore which processes could be behind higher order interactions in species-rich consumer-resource communities [4, 5].

### Holling Type II

Throughout this note, let us remark from the outset that we use predator-prey and consumer-resource as fully interchangeable terms, respectively.

#### A consumer on a single resource

In the most simple case, consumers can attack resource items, and then spend some time handling and/or digesting them, until they get ready to attack and capture new resources again. Therefore, feeding dynamics can be simply described by the following two processes:

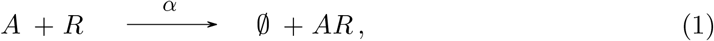

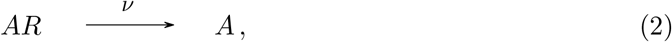

where individual consumers *A* encounter resource items *R* at a rate *α* creating a pair or “compound” denoted as *A*, i.e., a handling predator, which “relaxes back” into a free consumer at rate *v*. Although here space is not explicitly considered, the empty set in Eq. (1) emphasizes that the actual interaction may involve the jump of the predator onto the prey, which would leave empty space behind. In addition, in this derivation, we consider that resource density is kept constant, at the same resource level, which we noted 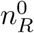 Since no reaction leading to consumer depletion is assumed, the total number of consumers, 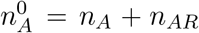 is also kept constant. In analogy to chemical kinetics, we call these conditions of constant population for both prey and predators *chemostatic consitions* [18]. Consumers can be then in two behavioral stages, either searching new resources or handling them, *A* and *AR*, respectively. At any given time, since the total number of consumers is constant, we can characterize the configuration of the system only by the number of searching consumers, *n*_*A*_, free and ready to attack new resources. If we assume that we can pack at most *N* resources in an given area where consumers *move randomly around*, their effective attack rate will be damped by the actual density of resources (which will be denoted as 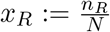) in that area. This common assumption is usually called true *mass action* [18, 19].

In this context, we can dynamically model the feeding process by using one-step stochastic processes in continuous time [17]. This formalism allows to express the temporal evolution of the probability of finding *n*_*A*_ free consumers at time *t, P* (*n*_*A*_, *t*) through a master equation, which can be solved for the steady state. In this case, at stationarity, *n*_*A*_ will follow a binomial distribution with parameters 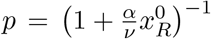 and 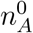(see Box I, Fig. 1 and Supplementary Information for details), where 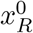 is the resource density. Accordingly, the average number of free consumers at stationarity will be given by 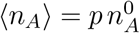. The average rate of resource depletion caused by the total population of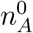 predators at stationarity will be given by 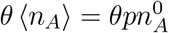, where we defined 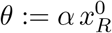. When this rate is expressed on a per capita basis, with respect to the total population of predators, it gives rise to a per capita averaged feeding rate, the Holling Type II functional response:

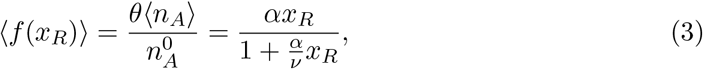

where we have finally dropped the superscript 0 on resource densities. This emphasizes that, while 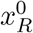 was a parameter throughout our derivation, it can be regarded as a true dynamic variable when Eq. (3) is used in population-level models. We will take this prescription also to write the final expressions of the functional responses throughout all the derivations that follow.

**Figure 1:**
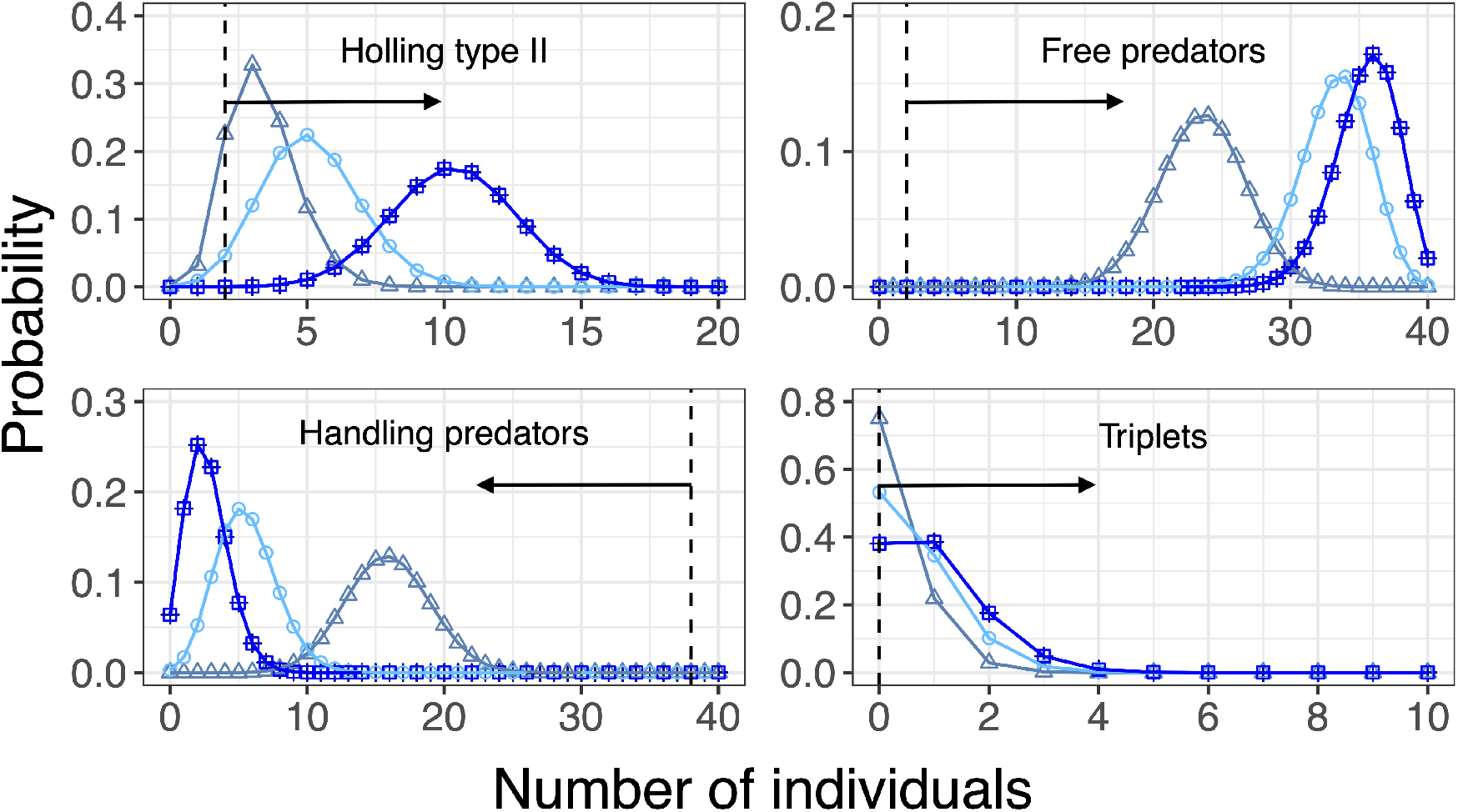
Probability distributions for: (i) free predators (top left panel) in the Holling type II case; (ii) free predators (top right), handling predators (bottom left), and triplet compounds (bottom right) in the case of predator interference dynamics (for simplicity, we considered the case *χ*_*AA*_ = *η*_*AA*_ = 0). In this figure we compare the numerical integration of the master equation (triangles, circles, and squares) with the asymptotic distribution (crosses) given by Eq. (S14) for the Holling (top left) case, and analytical solutions for the interference dynamics case (remaining panels). The integration of the master equation was carried out up to *t* = 3.06 (triangles), *t* = 8.16 (circles), and *t* = 50 (squares, Holling type II case) or *t* = 15 (squares, interference dynamics panels). The integration of the master equation shows a perfect agreement with the analytical, equilibrium distributions in the limit of *t* tending to *∞*. Parameters for Holling type II (top left) panel are: 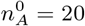 individual consumers (distributions’ support goes up to that value), with attack rate *α* = 2.5, handling rate *v* = 1 feeding on a (fixed) total resource density 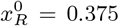. The master equation integration started with *n*_*A*_ = 2 free predators at *t* = 0 (dashed vertical line). Parameters for interference dynamics (top right and bottom) panels are: 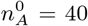 individual consumers (this is the support of free and handling predators distributions), with attack rate *α* = 2.5, handling rate *v* = 10, interference rates *χ* = 100 and *η* = 1, feeding on a (fixed) total resource density 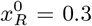. The support of the triplets’ distribution goes up to *n*_*ARA*_ = 20 in this example (this limit is not shown in the bottom right panel to ease visualization). The master equation integration started with *n*_*A*_ = 2 free predators, *n*_*AR*_ = 38 handling predators, and zero triplet compounds at *t* = 0 (dashed vertical lines). In every panel, arrows show increasing time starting from the initial conditions just described.

Notice that the classic expression by Holling includes the handling time, *τ*, which is the average time it takes for an individual consumer to handle and digest a resource item. In our framework, this is 1*/v*, the average relaxation time associated to the first order kinetics represented by reaction (2).

In a similar way we can derive the classic Holing type III functional response from individual-based reactions of the form

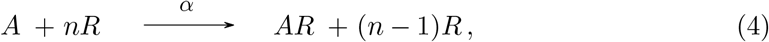

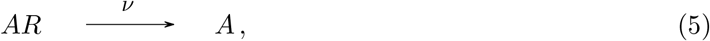

In an analogy with allosteric enzymatic reactions [10, 16], effective resource depletion rates are strongly affected by the density of the resources, which can be regarded as resource items and consumers forming higher order compounds. Under the same assumptions as before, at stationarity, this feeding mechanism gives rise to the same binomial distribution for the number of free consumers, but with a slight different definition of the *θ* parameter 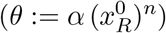,which end up determining an average feeding rate of the form:

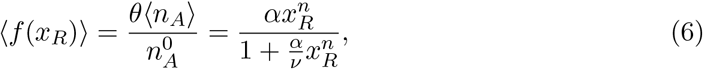

where *n* is the so called *Hill exponent*, the rest of parameters have the same meaning as in 3.

#### A consumer on multiple resources

We can ask now what is the correct form of the functional response when a Holling Type II consumer feeds on multiple resources. We will show that the answer to this question is not unique. Let us first consider that a consumer individual *A* can feed on a range of *S* alternative resources, *R*_*i*_ with *i* = 1, …, *S*. In this situation, the dynamics can be summarized by 2 *S* individual feeding reactions:

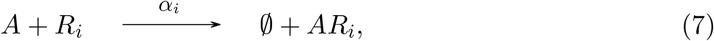

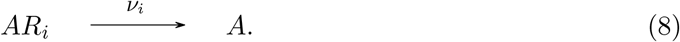

The first one takes into account resource preferences as involves distinct attack rates (*α*_*i*_) for the different resources, and the formation of handling consumers for each of the resources. Handling times (1*/v*_*i*_) are also characteristic for individual consumers feeding on each of the resource types. The total consumer population, 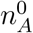 is divided in *S* + 1 classes, 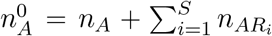. As before, we could also translate the reaction scheme (7)–(8) into transition probability rates between discrete configurations, and deploy the master equation driving the temporal evolution of the probability of finding a given discrete configuration of consumer types, 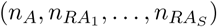, at a given time *t*, to determine the stationary state. This calculation is not straightforward and will be left for further work. Here, we simply present instead the deterministic rate equations emerging from this reaction scheme and calculate the average number of free consumers at stationarity, which we show leads to the average feeding rate per individual consumer when feeding on a given resource type.

For convenience, we first define *n*_*X*_ ≡ ⟨*n*_*X*_⟩, as continuous variable representing the different subpopulations of consumers: free consumers, *n*_*A*_, and handling consumers for each resource type, 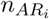. The rate of temporal change of these variables associated to the reaction scheme (7)–(8) can be written as:

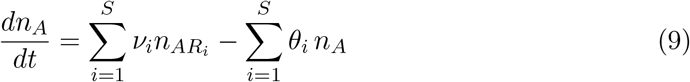

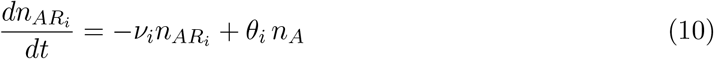

where the effective per capita attack rate of a free consumer at a given density level of the *i*-th resource type is a resource-specific constant parameter defined by:

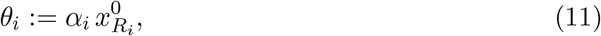

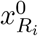 being the density of the *i*-th resource. The system represented by Eqs. (9)–(10) is a coupled system of *S* + 1 ordinary differential equations. Solving it for stationarity makes use that both consumers, 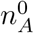, and all resource levels, 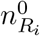, are kept constant, so all *θ*_*i*_ are constant parameters. Under these assumptions, at equilibrium we obtain the following linear system of equations:

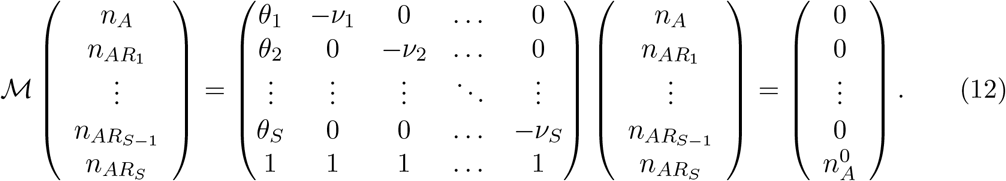

By inverting the matrix ℳ, we can find the distribution of the different consumer sub-populations at stationarity. Notice that 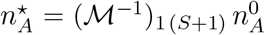, which immediately leads to:

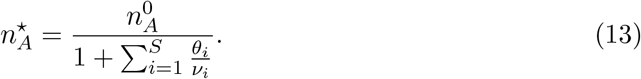

The subpopulation of free consumers 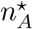is responsible for the total rate of resource loss at stationarity. In particular, resource *i* is consumed by ready-to-attack, free individuals at a per capita rate *θ*_*i*_. Therefore, total resource depletion of resource *i* per unit time is 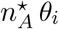 at stationarity, which, on a per capita basis, becomes the multi-resource Holling type II functional response of an average consumer feeding on a focal resource of the *i*-th type in the presence of alternative resource types:

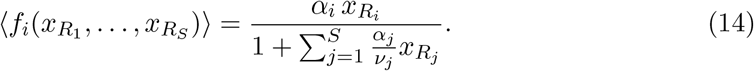

Both reaction schemes (1)–(2) and (7)–(8) assumed that the individual feeding process is not affected by the presence of either conspecifics (through interference), variable levels of resource density, nor different types of resources. In the multi-resource case, the feeding process is only controlled by the relative presence of the different resources, and the specific rates at which an individual consumer attacks and handles each resource type. In other words, individual attack rates (*α*_*i*_) and handling times (1/*v*_*i*_) for resource type *I* are the same regardless whether the individual feeds in total isolation or not, reacts differently to different levels of resource density, or when exposed to more than one resource type. If these parameter rates are not independent of all these factors, more complicated functional responses should be expected (see Suppl Mat). This approach can then be used to explore the origins of high-order interactions [5], which will end up determining different mathematical forms for the average per capita feeding rate of consumers when feeding on multiple resources. For instance, in the Supp Mat, we derive other possible multi-resource extensions of the Holling Type II functional response when more refined density dependence effects are present, and other mechanisms such as prey switching are considered.

In any case, although not always explicitly expressed, we should emphasize that both the single-Holling Type II functional response, and its multi-resource extensions are derived as (1) per capita asymptotic average feeding rates at stationarity, and under (2) chemostatic conditions, and (3) mass action dynamics.

### Beddington-DeAngelis: predator density-dependence

Two predators may eventually interfere by engaging in contests due to different reasons (fights for a territory, competition for resources, etc.). This interference behavior prevents them from resource search and consumption and, therefore, may potentially decrease population-averaged consumption rates. While the interference process lasts, the two consumers involved are entangled together. When the interference ends, the two consumers return simultaneously back again to the searching stage. This may lead to the formation of a “triplet”, made up by a handling predator-resource pair and a free predator, which breaks down into the pair and the free consumer, on average, after a finite amount of time. The disintegration of the the triplet can thus be modelled through a *relaxation* rate. This point is important because it has been overlooked in previous derivations of predator-dependent functional responses based on reaction schemes [13, 20]. These authors implicitly assume in their reaction schemes that two predators enter an interfering stage, simultaneously, at the moment they encounter each other, but relax back to the searching stage independently from each other, which is not consistent, and probably less realistic.

Here, we assume two types of interference processes. First, consumers can directly interfere while searching. Second, a searching consumer can encounter a consumer individual that is handling a resource item, entering in interaction, in the hope of obtaining perhaps a share of the resource. More complicated interference processes, including more complex behavioral effects, could be considered by adding other behavioral stages (consumers recovering from fights, consumer watching out for or hiding from top predators, etc.). The processes we considered here are minimal. They can be represented by the following reaction scheme:

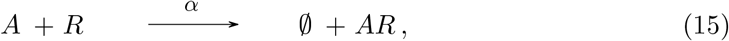

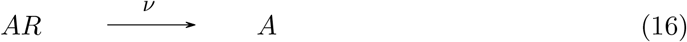

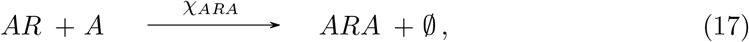

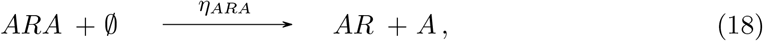

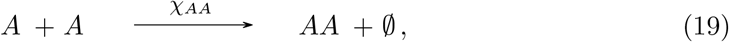

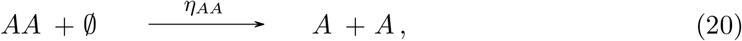

where we added the dynamics of pairs, *AA*, and what we call triplets, *ARA*. These are interference complexes, the first one formed by two searching individuals, *A* and *A*, and the second one formed by a searching individual, *A*, and a handling consumer, *AR*. Resource level, 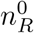, is assumed constant. As a consequence, the actual consumption rate of a free individual, *A*, is still given by 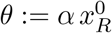. Since no process depleting the consumer population is considered, the total number of consumers, 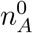, is also kept constant, and will be distributed among four different types. This adds a constraint to the dynamics: 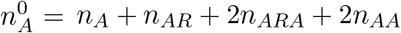. These types correspond to different behavioral stages: free consumers, *A*, handling/feeding consumers, *AR*, and interfering consumers engaged in either pairs, *AA*, or triplets, *ARA*. Once this distribution reaches steady state, there will be a steady number of free consumers, 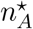, ready to search and attack new resources. This subpopulation will be directly responsible for total resource depletion at a constant steady-state rate, which will determine a population-averaged per capita feeding rate: the functional response.

Once again we could also translate the previous reaction scheme (15)–(20) into transition probability rates between discrete configurations (see Box I), and deploy the corresponding master equation to solve for the stationary distribution of behavioral types. The full derivation of such distribution is more convoluted and is beyond the scope of this note. It will be discussed in a forthcoming publication. Here, it is enough to consider rates equations for the continuous variables representing the different subpopulations of consumers, *n*_*A*_, *n*_*AR*_, *n*_*ARA*_, and *n*_*AA*_. Although this is an abuse of notation, for simplicity, we have used the same notation for both continuous deterministic and discrete stochastic variables. The context will make it clear. The value of 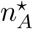 is defined by the steady-state that emerges from the deterministic rate equations associated to the reaction scheme (15)–(20), which reads:

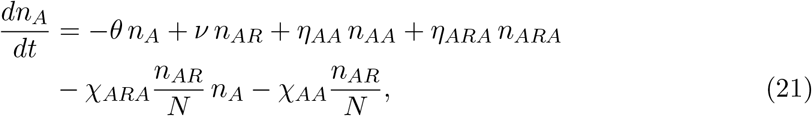

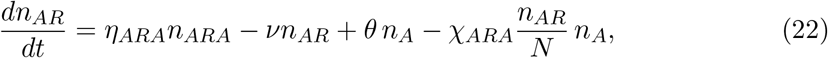

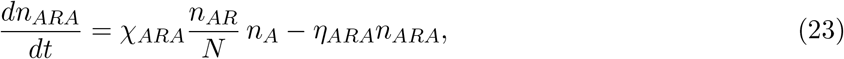

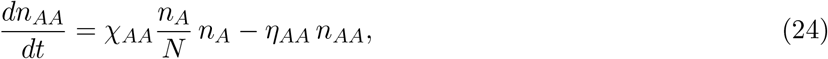

where *N* is again the number of resources that can be packed in the area of study.

By making the rate equations equal to zero, and using the constraint given by 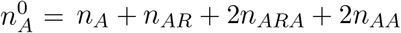, we obtain a value for *n*_*A*_ at steady-state:

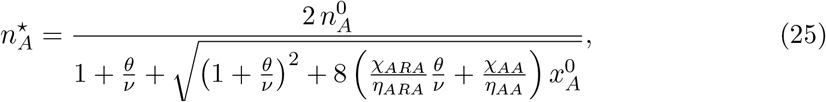

which leads to the expression of the population-averaged per capita feeding rate:

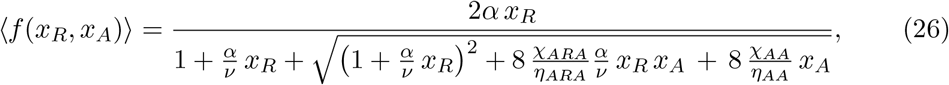

where we recall that we have dropped the 0 superscripts for simplicity and defined the densities 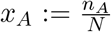 and 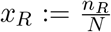. The new functional response derived by this methodology (Eq. (26)) leads, in certain limits, to the Beddington-DeAngelis functional response in its usual form. One possible way to recover Beddington-DeAngelis functional response from Eq. (26) is when there is only resource-mediated interference (*χ*_*AA*_ = 0), and attack and interference rates, relative to their corresponding relaxation rates, satisfy the conditions *α/v* ≪ 1 and *χ*_*ARA*_*/η*_*ARA*_ ≪ 1. This is a quite restrictive situation. In other limits, Beddington-DeAngelis functional response is not compatible with the reaction scheme proposed above.

As for the Holling type II functional response (see Box I), when interference is considered, the stationary distribution for the number of different behavioral types can always be obtained also through numerical integration of the master equation (see Box I and Supplementary Information). The temporal dependence of the distributions of behavioral types, calculated by numerical integration of the master equation, is provided in Fig. 1.

In future work, we will present further mathematical details on the derivation of these stationary distributions, their time-dependent behavior, and their potential use to infer feeding rates from experimental data both in transient and stationary situations.

Notice that Eq. (26) is the predator-dependent functional response that most consistently generalizes Holling Type II. If there is no interference (*χ*_*AA*_ = *χ*_*ARA*_ = 0), we recover Eq. (3). If there is only resource-mediated interference (*χ*_*AA*_ = 0), we recover the functional response we presented in [16]. This nested property is very useful for model selection and parameter inference. Previous derivations of the predator-dependent functional response [13, 20] clearly do not pass this check.

## Discussion

Functional responses can be regarded as a very early example in ecology of a general research agenda in complex systems science: how can processes at a lower level of aggregation —for instance, the individual level— be used to approximate the macroscopic dynamics of a system at a higher level of aggregation —for instance, the population or the community level? Methods to derive macroscopic equations from the elementary processes affecting individuals are continuously developed in statistical physics, ecology and beyond [21]. One well-established research line considers individuals as interacting particles and describes their interactions in terms of chemical kinetics [18]. The use of this approach in population, community ecology, and the dynamics of infectious diseases has become widespread [22, 23, 24, 25, 17, 26, 21, 27].

The three central inter-related messages of this paper are: (1) the link between individual processes and average feeding rates through stochastic processes, (2) the unavoidable implicit assumptions that this link entails for the derivation of the associated distributions and average quantities, and (3) the derivation of a new functional response, which can be regarded as the natural extension of the Holling type II (rather than the classic form from Beddington-De Angelis), when two types of predator interference are considered.

In addition, we have shown that reaction schemes represent a common framework to derive both single-resource functional responses and their multi-resource extensions [4, 5, 28, 29]. This approach has several advantages. First, the underlying hypotheses and particular assumptions leading to the different functional forms become evident. Second, through the use of stochastic processes in continuous time, not only expected average feeding rates but also their whole distribution across replicates can be calculated. This provides a natural link between individual stochastic descriptions of the feeding processes, and their population-averaged deterministic analogs [17]. And, third, the resulting functional responses are fully nested, which is a useful property in model selection contexts. However, it is important to emphasize also the limitations of the functional responses emerging from reaction schemes. First, they are population-level, per capita averages only asymptotically valid at stationarity. Second, they are based on the mass action assumption (individuals encounter resources and each other at random), and third, these derivations also rely on chemostatic conditions (resource and consumer population levels are kept constant while the steady state is reached). Stationarity (1), mass action (2), and chemostatic conditions (3) are three implicit assumptions underlying classic derivations of functional responses that have been sometimes overlooked.

Chemostatic and stationary conditions are related. Stationarity should be reached fast enough before any extra processes affecting resource and consumer population levels becomes important. Common use of functional responses of this type in population dynamic models requires in general a strong time scale separation between a fast (behavioral) time scale controlling feeding dynamics, and a slow (demographic) time scale characterizing processes such as death, migration and reproduction. This strong time scale separation allows for feeding dynamics to reach the steady-state characterized by average consumption rates, which can be then introduced in the population-level model. This separation of time scales is emphasized in recent derivations of classic consumer-resource models from individual processes, such as the Rosenzweig-MacArthur and Beddington-DeAngelis consumer-resource models, which consider a consumer species feeding on a logistically growing resource [16, 30, 31]. The use of these population models relies implicitly on this strong time-scale separation assumption. Population equations with classical functional responses are thus effective models in the sense that they encapsulate the whole complexity of a low-level behavioral process occurring at a rapid time scale in a single per capita average macroscopic feeding rate, which in turn depends on certain densities and effective parameters. For instance, a time-scale separation argument allows to recover the classic Rosenzweig-MacArthur consumer-resource model from slightly different feeding and population processes. However, the precise interpretation of functional-response effective parameters differs in each case [16, 30, 31].

Interactions rates are governed by the rate at which individuals encounter, and this, in turn, results from the way individuals move, when, for instance, they search for food. A rich research area focuses on both measuring and theoretically deriving optimal movement distributions of foraging animals depending on total food density and its distribution over an area [32, 33, 34]. In the search for functional responses, a recent study shows that different functional forms emerge when individuals do not mover randomly but following more realistic trajectories, such as Levy walks [35]. Advances in technology (optics, data compression, computation capacity, robotics) have made it increasingly feasible to automate behavioral assays in the laboratory and thereby collect more animal data with minimal human intervention [36]. Feeding experiments can potentially benefit a lot from these novel methods in conjunction with the ideas presented in this note. More research is needed to understand to which extent the distribution of behavioral types could allow a better estimate of the functional response parameters. This approach would require the ability to recognize distinct predator behavioral stages. Predators in different behavioral feeding stages may move in recognizable, distinct ways. In fact, the segmentation of moving trajectories into distinct behavioural stages is becoming more and more accessible due to novel machine learning and artificial intelligence methods [37, 38].

In recent years, functional responses have been addressed by using different stochastic approaches. Van der Meer and Smallegange [20] use a stochastic analysis that differs from ours because it focus only on the stationary distribution of a discrete-time model. Billiard et al. [39] make use of general renewal processes to estimate the average and the variance of feeding rates as an approximation when the number of consumers is large. Here, we emphasize that chemical-like reactions schemes provide a straightforward natural link between deterministic and stochastic dynamics because they can lead to both deterministic rate equations for average densities, and one-step stochastic master equations in continuous time for the whole exact probability distribution of the different behavioral types. We highlight that this property has a huge potential for parameter inference and model selection in the context of feeding experiments, particularly, when the number of prey and predators is low.

Finally, several derivations of the Beddington-DeAngelis functional response abound in the literature [13, 20, 40, 41], which describe predator’s interference at both the individual and the population level. However, the derivation we present in this note, resulting in Eq. (26), is the only one that considers the same three common assumptions as the classic Holling Type II functional response, and should be then regarded as its natural generalization, under both direct and resource mediated interference. It is only under very restricted conditions that this expression collapses into the classic Beddington-DeAngelis functional response [16], and, only within these strict conditions, we can safely say that the classic expression approximates the true per capita average feeding rate of interfering consumers. According to Eq. (26), when consumer interference for resources (formation of triplets) is strong, this is, dominates over the rest of the feeding processes, then total resource depletion, i.e., 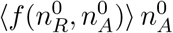, scales as the square root of the product of prey and predator densities. Interestingly, this square root scaling prediction has been found in recent empirical estimates of total predation rates from macroecological food-web data across many species and orders of magnitude in biomass [42]. The use of the insights we have gained with the work presented here to better understand food webs relies on the three assumptions mentioned before. In particular, systems should be considered at stationarity along with a clear separation of behavioral and demographic temporal scales. In the context of the current ecological crises affecting ecosystems at local and global scales, urgent work is needed to understand perturbed and out-of-equilibrium systems across scales. In this direction, preliminary work is already begin conducted [43, 44].

### Box I

**Steady-state distributions of behavioral types**

**Holling Type II** feeding dynamics can be simply described by two discrete events: (1)the attack of resources, this is, an event that decreases the number of searching predators by one, and (2) the handling process, an event by which a handling predator relax back into the searching stage, and, as a consequence, the number of searching predators increases by one. This type of stochastic processes are known as birth-death or one-step stochastic processes. The master equation for one-step one-dimensional stochastic processes can be written as [18]:

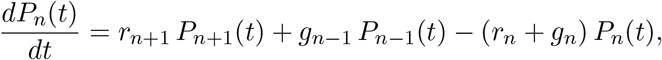

where *n* is a non-negative integer number in population contexts, and *P*_*n*_(*t*) is the probability for the system to have *n* individuals at time *t*. Notice that *r*_0_ = 0, since the population cannot decrease from zero. In this situation, the last equation is valid only as long as *n >* 0, and a particular equation determines *P*_0_(*t*):

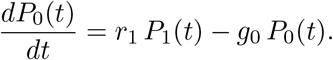

The feeding process described by Eqs. (1)–(2) is governed by the transition probability rates:

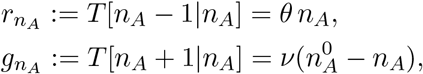

where *θ* is the effective per capita attack rate of a free consumer at a given level of resource density, i.e., here a constant parameter defined by 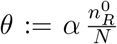, and 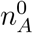 is the overall number of predators (constant as well). These rates control the temporal evolution of the probability of finding *n*_*A*_ free consumers at time *t, P* (*n*_*A*_, *t*).

As *t* → ∞, the distribution *P* (*n*_*A*_, *t*) may tend to a stationarity. Let *P* (*n*_*A*_) be this stationary distribution. We show in the Supplement that

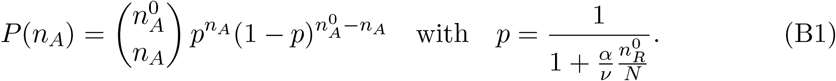

Therefore, we obtain that the stochastic variable *n*_*A*_ follows the binomial distribution at stationarity for Holling Type II feeding dynamics.

## Supplementary Material

### S.1. Stationary distribution of predator behavioral types

#### S.1.1. Holling Type II feeding dynamics

As mentioned in the main text, Holling Type II feeding dynamics is described as a birth-death stochastic process (also known as one-step process), because elemental changes either increase or decrease population numbers by one. These two events are: (i) predators attack resources, so the number of searching (free) predators decreases by one while attacking the prey, and (ii) handling predators abandon the resource and come back to the searching stage, so the number of searching predators increases by one.

For a general one-step one-dimensional stochastic process (i.e., a process that focuses on a single stochastic variable, in this case, the population number of free predators in the system), the general expression of the master equation is [18]:

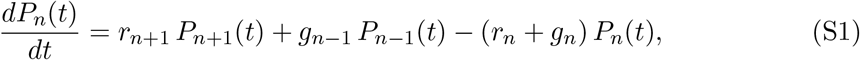

where, in general, *n* is an integer number, and *P*_*n*_(*t*) denotes the probability of observing the value *n* at (continuous) time *t*. The temporal dynamics *n*(*t*) is described by the fraction of times a particular value of *n* is observed at time *t* over a large number of stochastic realizations of the process, i.e., by the probability *P*_*n*_(*t*). The dynamics is mathematically defined once the transition probability death and growth rates (*r*_*n*_ and *g*_*n*_, respectively) are specified as functions of *n*.

For convenience, we re-write the master equation as:

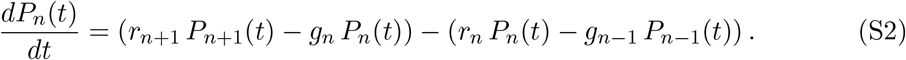

In population contexts, *n* should be a positive integer representing the number of individuals in the population, and then *P*_*n*_(*t*) is the probability for the system to have *n* individuals at time *t*. The death rate must satisfy *r*_0_ = 0, because the population number cannot decrease from zero. In this situation, Eq. (S2) is valid only for all probabilities as long as *n >* 0, and *P*_0_(*t*) is determined by

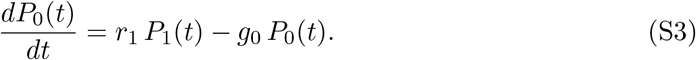

As *t* → ∞, the distribution *P*_*n*_(*t*) may reach a steady-state distribution which is independent of *t*. Let *P* (*n*) = *P*_*n*_ be this stationary distribution. By definition, this probability distribution should make all right-hand sides of Eqs. (S2)–(S3) for all *n* vanish, so the (infinite) set of coupled differential equations reaches an equilibrium point. A sufficient condition for stationarity is the so-called detailed balance condition,

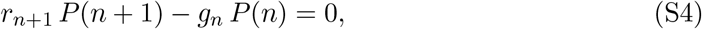

which has to be satisfied for all *n* ≥ 0. Due to the telescopic form of (S4), the two terms in the right hand side of (S2) vanish independently. If we find a solution for the detailed balance condition, then we will have found the stationary distribution.

For *n* = 0, Eq. (S4) is basically the condition that equals to zero the right-hand side of Eq. (S3). Then it follows that the first two values of the steady-state probability distribution, *P* (0) and *P* (1), should be in the right proportion:

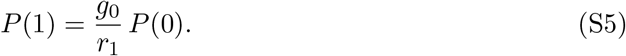

Eq. (S5) for *n* = 1 yields

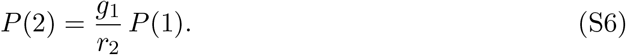

If we repeat this process, recurrently, for *n* = 2 and so on, we obtain the general expression for *P* (*n*):

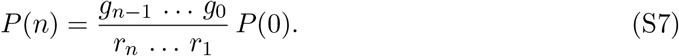

This simple relation gives the stationary distribution up to a normalization constant *P* (0).

Now we particularize the distribution for Holling type II feeding dynamics defined by the transition probability rates

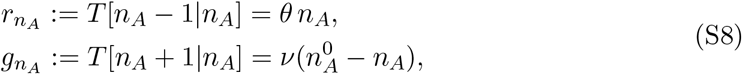

where *v* is the per-capita probability rate of *AR* handling predator disintegration, and 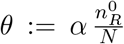is the product of the rate *α* of *AR* handling predator formation times the number of resources 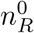 relative to the total number of sites *N*. Therefore, *θ* is a constant because the resource level 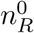 is assumed to be constant.

If we use then the probability rates (S8) in Eq. (S7) —where *n* is now the number of searching predators, *n*_*A*_— to determine the stationary distribution *P* (*n*_*A*_) of the number of free predators, we obtain

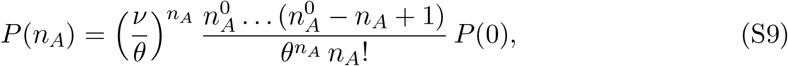

which can be written as

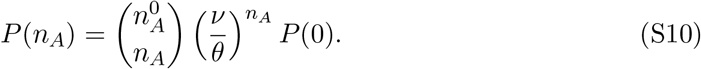

This computes the distribution up to a normalization constant *P* (0), which can be obtained by the normalization condition, 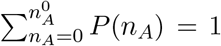. This condition, together with (S7) determines *P* (0) as

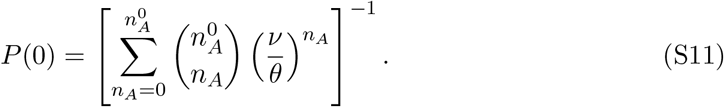

Using the binomial expansion we can write

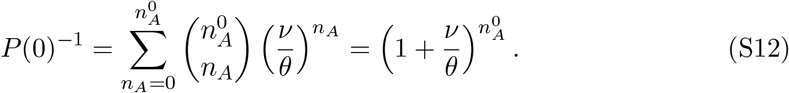

Therefore,

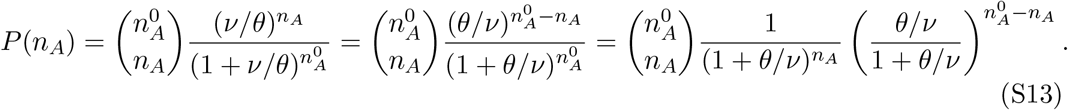

Finally, we obtain that the stochastic variable *n*_*A*_ follows the binomial distribution 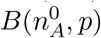 at stationarity, with probability distribution function given by

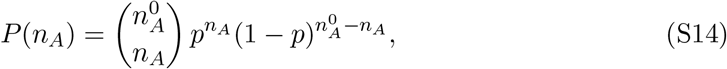

with parameters 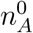 and

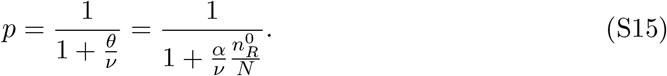

### S.2. Multi-resource Holling Type II

#### S.2.1. Density effects through resource interactions

Consumer efficiency on a focal resource can be intrinsically affected not only by the level of the focal resource, as in Holling type II functional response, but also by the presence of alternative resources. In order to take this observation into account, the chemical scheme in reactions (7)–(8) of the main text should be extended, and this can be done in different ways. Consider, for instance, a non-linear increase (or decrease) of feeding efficiency in response to both the same and alternative resource types. This would correspond to the multispecies version of Holling type III (where a Hill exponent *n* = 2 has been assumed as the power to which resource density is raised in the functional response):

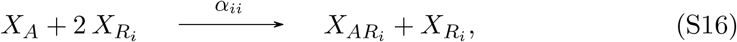

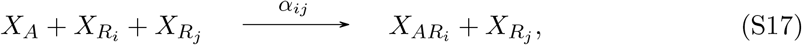

where *α*_*ij*_ are reaction rates taking into account the effect of the density of the same (*i* = *j*) or alternative resources (*i* ≠ *j*) on the formation of handling consumers of the focal resource *i*. When comparing rates *α*_*ii*_ and *α*_*ij*_, we can see whether the feeding rate on *R*_*i*_ is enhanced (or damped) by the presence of an alternative resource *R*_*j*_ more than it is by the presence of the same type of resource *R*_*i*_. Notice that, in this case, the full reaction scheme of the individual feeding processes will be reaction (8) of the main text together with reactions (S16)–(S17) defined above, both of which replace channel (7), see main text. The per capita consumption rate of a free/searching individual *X*_*A*_ feeding on a focal resource *i* has now two terms:

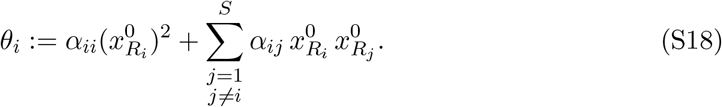

Interestingly, the same rate equations as the ones used for the previous derivation of the multi-resource Holling Type II funcional response —see Eqs. (9) and (10) of the main text— apply here but now using this new definition of *θ*_*i*_, which is still a constant parameter that depends on resource type. Therefore, the same reasoning that led to Eq. (14) in the main text, now yields the following general multi-resource functional response:

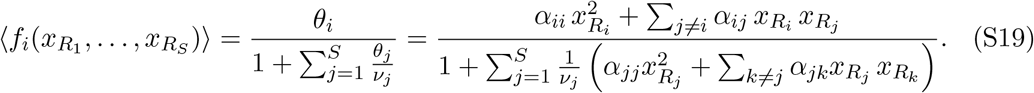

Observe that, if we consider the reaction given by Eq. (7) (main text) together with (8) (main text) and Eqs. (S16)–(S17), i.e., we allow for processes in which the predator encounters a single resource item of class *i* and processes in which predator efficiency is affected by the presence of two resources from equal or different classes, the same derivation of the multi-resource functional response (S19) will hold, in this case yielding a functional response that combines Hill exponents *n* = 1 and *n* = 2. The only difference is that *θ*_*i*_ in Eq. (S18) should be replaced by

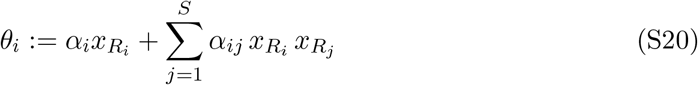

when substituted into

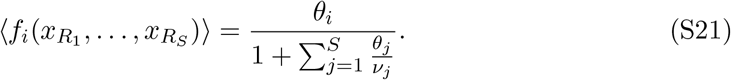

#### S.2.2. Prey Switching

Here we consider that individuals *X*_*A*_ can feed on a range of *S* alternative resources, 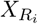 with *i* = 1, …, *S*. In addition, handling consumers, when they encounter another resource type, can release the one they have been handling and switch to the new one (at rate *η*_*ij*_) —prey switching.

In this situation, the dynamics can be summarized by 4×*S* individual feeding reactions:

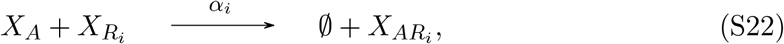

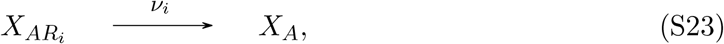

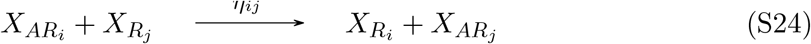

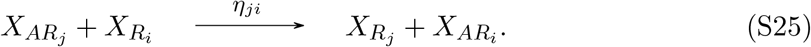

When we consider the full feeding hypothesis given by the reaction scheme in (S22)– (S25), the associated rate equations read

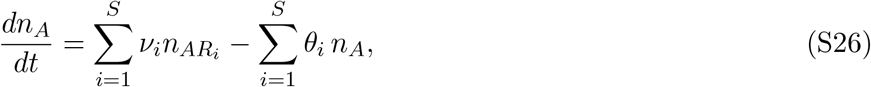

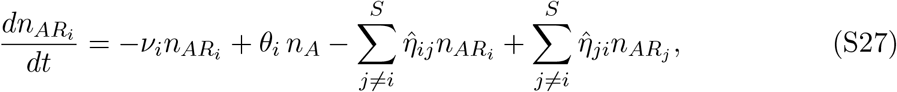

where we have defined

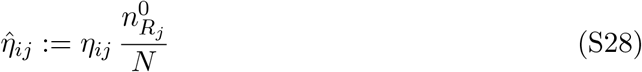

and *θ*_*i*_ is still given by (see Eq. (8) in the main text)

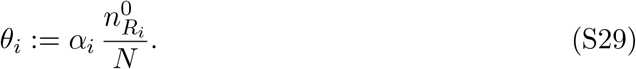

This is a coupled system of *S* + 1 ODEs. Solving it for stationarity makes use that both consumers, 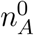, and all resource levels, 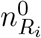, are kept constant, so all 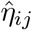 and *θ*_*i*_ can be considered as constant parameters. Under these assumptions, we obtain a linear system of equations, analogous to Eq. (12) of the main text, for the different subtypes of consumers at stationarity, 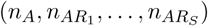, where now the ℳ matrix reads:

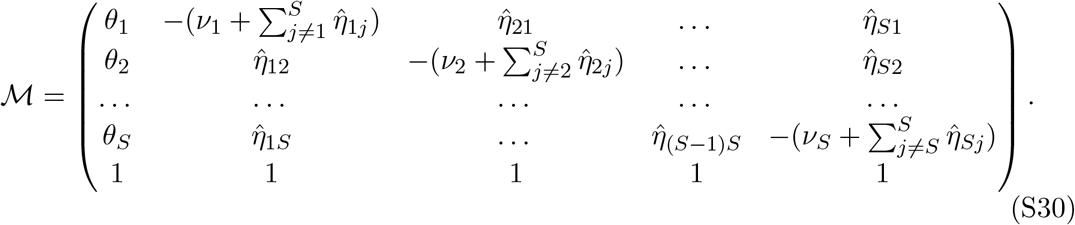

Again 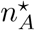 can be calculated in terms of a single element of the inverse matrix, ℳ*−*1, this is, 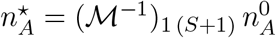, which leads the following functional response:

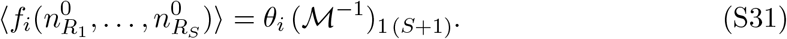

This involves the computation of an inverse matrix to obtain the average functional multi-resource Holling type II rate.

These per capita average rates of consumers feeding on a focal resource *i* results from specific assumptions that consider the type of higher order interactions (HOI) clearly specified by a full reaction scheme (S16)–(S17) and (S22)–(S25). Different types of HOI can be considered. In general, different reaction schemes lead to a different expressions for the functional response of consumers feeding on multiple resources. These two examples of average feeding rates in a multi-resource context are meant to emphasize that specific expressions of the functional response are contingent to the particular hypotheses assumed by a given reaction scheme. Although it is true that two different hypotheses on the individual feeding mechanism may lead the the same population-averaged per capita feeding rate of consumers [5], in general, parameter interpretation will be completely different, and this coincidence would not be the rule but rather the exception.

